# Stacked Waves and Qualitative Analysis of an Oncolytic Virus-Immune Model

**DOI:** 10.1101/2025.09.04.674336

**Authors:** Negar Mohammadnejad, Thomas Hillen

## Abstract

Oncolytic virotherapy (OVT) represents an innovative and promising therapeutic method for cancer treatment. This approach involves the introduction of oncolytic viruses into the patient, which are engineered to selectively target and lyse tumor cells. Based on previous mathematical modelling of oncolytic viruses, we consider a mathematical model that describes the intricate interactions between the oncolytic virus, cancer cell populations, and the immune system. Our study includes a detailed qualitative and quantitative analysis of the model to explain why, despite their promise, oncolytic viruses alone rarely lead to complete and lasting regression of established tumors in vivo. We use parameter sensitivity analysis to support our findings. Furthermore, we consider the spatial version of the model in the form of a system of reaction-diffusion equations. A travelling wave analysis shows an unexpected phenomenon. The solution components do not necessarily evolve into a single traveling front; rather, they develop into stacked fronts, where each front propagates at a different speed. We give explicit formulas for these different invasion speeds, confirm those through numerical simulations, and discuss their significance for OVT.

## 1 Introduction

Oncolytic virotherapy (OVT) is an emerging cancer treatment in which oncolytic viruses are used to eradicate tumor cells. One of the most remarkable properties of OVs is their ability to selectively infect, replicate, and propagate within tumors (Engeland et al., 2020). The first genetically engineered OV, a herpes simplex virus 1 (HSV-1) mutant with deficient thymidine kinase, was developed in 1991 and used to treat malignant glioma in nude mice. These viruses selectively infect and destroy cancer cells, often engaging dendritic cells to enhance immune-mediated antitumor responses (Kim et al., 2015). Although oncolytic viruses (OVs) have advanced into clinical trials, with T-VEC receiving FDA approval for melanoma, significant challenges remain, including limited viral infection efficiency and rapid immune clearance, which must be addressed to fully realize their therapeutic potential (Mahdie et al., 2022).

The immune system plays a pivotal role in OV–cancer interactions. The immune response was once considered as a virus replication inhibitor and a limiting factor in virus-cancer interactions. However, the interactions of cancer, virus and immune response are more complex. The immune system recognizes and attacks the tumour after the viral infection enhances the visibility of the tumour to the immune system. Moreover, the release of tumor antigens into the microenvironment, triggered by virus-induced lysis of cancer cells, helps reverse immune suppression which is often present in a tumor tissue.

### 1.1 Paper Outline

In this work, in Section 2, we start with constructing a cancer-virus-immune model to analyze the complex dynamics that might arise during OV therapy. Inspired by the work of Baabdulla and Hillen (2024) and Al-Tuwairqi et al. (2020), we incorporate the population of immune cells into the OV model studied by Baabdulla and Hillen (2024). We investigate the fundamental interactions of immune response with cancer and with the oncolytic virus. In Section 3, We will first consider the space independent model without the diffusion terms and we conduct a qualitative analysis of the model and its steady states. We examine the stability of equilibrium points and we show how the Hopf bifurcation, known from the OV-model of Baabdulla and Hillen (2024), is affected by the presence of the immune response. We find that the immune response, if strong enough, will inhibit oscillations. Next, we perform a comprehensive sensitivity analysis of all model parameters. This analysis highlights that the growth and decay parameters of the immune response are most sensitive to the treatment outcome.

In section 4, we put the OV-immune model into a spacial context and include diffusion terms into the model equations. Of particular interest are the invasion speeds of the model. A travelling wave analysis shows an unexpected phenomenon. The solution components do not necessarily evolve into a single traveling front; rather, they develop into stacked fronts, where each front propagates at a different speed. We see that the faster components advance ahead while slower ones trail behind. Specficially, we observe a first invasion front as cancer invades the healthy tissue. This is followed by a viral wave front that catches up with the cancer wave. Finally, an immune response wave front follows the previous two and might or might not catch up. Instead of a single coherent wave, the system exhibits a layered structure of stacked traveling fronts. We close with a Conclusion Section 5.

Before presenting our model in Section 2, we first provide a brief overview of some of the previous works on OVT modelling and stacked invasion fronts.

### 1.2 Previous Modeling of OV Therapy

One of the first OVT models was introduced by Wodarz (2001). He explored how virotherapy could be optimized to achieve maximal therapeutic effects. Wodarz also examined three main mechanisms of action: direct viral killing of tumor cells, immune responses against the virus that indirectly aid therapy, and the induction of tumor-specific immune responses triggered by the infection. The works Wodarz and Komarova (2009), and Komarova and Wodarz (2010) on oncolytic virotherapy have served as a foundational framework upon which more complex models have been constructed. In 2009, they started with presenting a broader modeling framework that accounted for different modes of virus spread (fast vs. slow) and tumor growth kinetics (Wodarz and Komarova, 2009). Their key insight was that the dynamics of OV therapy can fall into two qualitatively distinct regimes, the fast spread and the slow spread. They continued their study on OVT dynamics in the following year. In Komarova and Wodarz (2010), they developed a general system of ODEs that is based on the biologically motivated properties of tumor growth and viral spread. They analyzed how varying these assumptions would alter the system dynamics and defined conditions under which virus-mediated tumor eradication would be possible. Their work also underscored the importance of model robustness and cautioned that many classical modeling results may be artifacts of arbitrary mathematical choices. Tian (2011) introduced a model comprising three variables, the uninfected and infected cancer cells, and the virus. His findings indicated that virotherapy success strongly depends on the burst size of the oncolytic virus. A 3D spatio-temporal model was developed by Pooladvand et al. (2021). They studied the dynamics of cancer-virus interactions, specifically focusing on adenovirus therapy within a solid tumor. In their study, by linking PDE modeling with bifurcation analysis, they revealed deep insights into why virotherapy often falls short as a standalone treatment, and underlined the limitations of virotherapy as a monotherapy. Their work inspired the model by Baabdulla and Hillen (2024), who developed a diffusion model of OVT and extensively studied the temporal and spatial interactions of the virus, and the cancer cells, but without considering the effect of the immune system. In the absence of OVs, tumors typically establish an immunosuppressive microenvironment that suppresses the body’s immune response. The introduction of oncolytic viruses, however, triggers pro-inflammatory signaling within the tumor. There has been extensive research on the role of the immune system in the cancer-virus interactions, primarily emphasized on its role in mediating these interactions. For instance, Eftimie et al. (2011) developed an ODE-based model with two immune compartments—lymphoid and peripheral—to capture the dynamics among uninfected and infected tumor cells, memory and effector immune cells, and two types of viruses: adenovirus (Ad) and vesicular stomatitis virus (VSV). Their model successfully replicated experimental tumor growth and immune response patterns, enabling an exploration of conditions required for sustained tumor elimination. Their analysis centered on complex behaviors such as multistability and multi-instability, revealing that persistent viral presence was associated with the latter. They also found that viral persistence alone was insufficient for tumor clearance without a concurrent anti-tumor immune response, underscoring the necessity of combining viral and immune-mediated effects for effective therapy. In another study, investigated the distinct contributions of innate and adaptive immunity in OVT by considering the adaptive immune cells as tumor- or virus-specific. They noted that the innate immune system often acts too rapidly, clearing the virus before it can adequately infect tumor cells and activate a strong anti-tumor immune response. Consequently, OVT by itself frequently fails to eliminate tumors. To address this, they modeled a combination therapy involving OVT and a PD-1/PD-L1 checkpoint inhibitor. This approach mitigated T-cell exhaustion, enhanced immune-mediated tumor destruction, and reduced the required viral infectivity threshold for successful treatment. Their findings emphasized the importance of optimal treatment scheduling and dosing, noting that administering a second viral dose too soon could inadvertently redirect the immune response toward the virus rather than the tumor, ultimately compromising therapeutic efficacy.

### 1.3 Invasion Fronts and Stacked Waves

In 1975 and 1978, the contributions of Aronson and Weinberger (1975, 1978) established the concept of spreading speed in single-species models with monotone nonlinearities, showing that this speed coincides with the minimal velocity of traveling waves. Building on this foundation, Weinberger (1982) later developed a powerful recursive approach to characterize spreading speeds in single-species systems, providing a versatile tool for analyzing wave propagation in such models. Later the definition of stacked waves was developed. Feinberg and Terman (1991) showed that if there are multiple distinct, locally stable euilibrium points and all of the equilibria are nondegenerate, then there must exists a wave train which connects them. Later Roquejoffre et al. (1996) discussed the convergence of the traveling wave fronts of a monotone parabolic system toward stacked families of waves. The existence of stacked waves has been studied by different authors. Iida et al. (2011) showed under certain conditions a cooperative systems with equal diffusion coefficients propagate at the same speed or they develop into stacked fronts where each front propagates at a different speed than the others. In the work by Hamelin et al. (2022), a coupled reaction-diffusion model is used to describe pathogen spread in genetically diverse plant populations. The interaction of pathogen genotype and plant species resistance let to the emergence of two successive invasion fronts. The first one dominated by the wild type pathogen and the susceptible plant population and the latter wave dominated by the resistant plant population and the resistance-breaking pathogen. Du and Wu (2018) showed in the weak–strong competition case, two invasive species can spread successfully but at different speeds, leading to spatial segregation of their populations. They studied two-component reaction–diffusion systems and proved that spreading occurs with well-defined speeds, showing the intriguing possibility that a single solution can generate two distinct fronts moving at different speeds. In the work done by Ducrot et al. (2019) on a two-component reaction–diffusion system, they proved that a single solution can generate two distinct fronts moving at different speeds. Liu et al. (2021) analyzed a three-species competition-diffusion system spreading at different speeds and showed that the spreading speed of the slowest species is dependent on the spreading speeds of the two faster one.

## 2 The mathematical model

Our model for immune response to oncolytic virotherapy is based on a standard oncolytic virus model as analyzed recently in Baabdulla and Hillen (2024). We want to incorporate the population of effector immune cells *Y* into their model, so our proposed model is given by

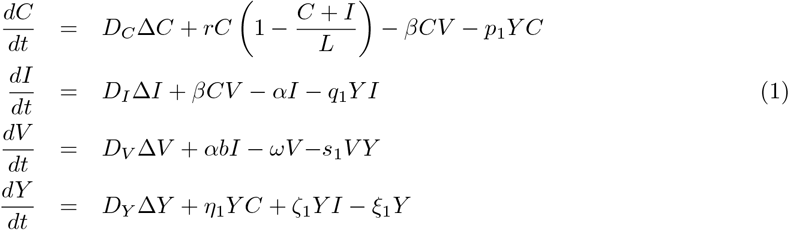

where *C*(*x, t*), *I*(*x, t*), *V* (*x, t*), and *Y* (*x, t*) represent the populations of uninfected cancer cells, infected cancer cells, free virus particles, and the immune cells respectively. Their corresponding diffusion coefficients are denoted by *D*_*C*_, *D*_*I*_, *D*_*V*_, and *D*_*Y*_. We have the Laplace operator Δ as the sum of all second order derivatives. The second term in equation (1) describes the logistic growth of cancer cells in the absence of therapy at the rate *r* with carrying capacity *L*. The interactions between tumor cells and virus particles are modeled by the mass action term *βCV* where cancer cells become infected by the virus at a rate *β*. The second term in this equation accounts for the increase in the infected cells. The lysis of these infected cells is represented by the third term at the rate *α*. In the third equation, *b* represents the burst size of the virus in infected cells, while *ω* denotes the viral clearance rate. The effect of the immune response on cancer cells, infected cancer cells, and the virus is modeled through standard mass action terms with rates *p*_1_, *q*_1_, and *s*_1_, respectively. The last equation of this system describes the stimulation of the immune system by both uninfected and infected cancer cells with rates *η*_1_ and *ζ*_1_, respectively. The immune cell are cleared at rate *ξ*_1_ as denoted in the last term.

We consider the system (1) with the initial conditions:

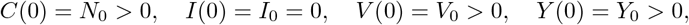

and homogeneous Nuemann boundary conditions on *∂*Ω

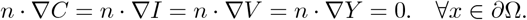

### 2.1 Non-dimensionalization

To simplify our analysis, we apply the non-dimensionalization method by considering

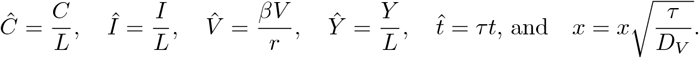

After dropping the hats, we obtain the non-dimensionalized system:

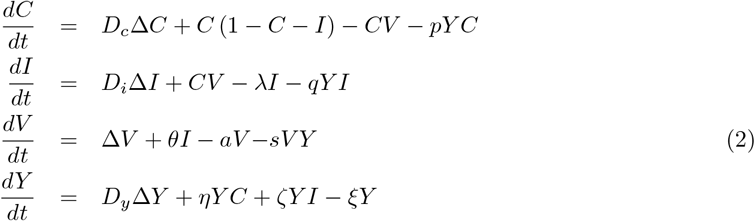

where

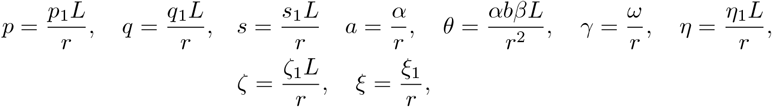

and

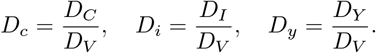

The parameter values before and after non-dimentionalization are given in Tables 2 and 3 respectively. The parameter values used in our simulations are chosen from different sources. Based on the work by Lodish (2013), Pooladvand et al. (2021) considered the carrying capacity of a solid tumor of radius 1 mm is about *L* = 10^6^ cells per mm^3^. We also chose this value as the initial density of uninfected tumor cells. In the experiments carried out by Kim et al. (2006) on adenovirus in the glioblastoma U343 cell line, the tumor growth rate was estimated to be approximately 0.3 per day. We will use this value in our analysis. We used the same parameter values for *L, β, ω*, and *b*, together with the initial condition for the adenovirus load (*V*_0_ = 1.9 *×* 10^10^ virions per mm^3^), as reported in the model of Baabdulla and Hillen (2024), where these parameters were associated with a reovirus and applied in the context of breast cancer. The remaining parameters used in our model are estimates of parameters in the model by Al-Tuwairqi et al. (2020). The parameters concerning the virus in their model for the glioma treatment were derived from the study by Friedman et al. (2006), which focused on the mutant herpes simplex virus 1 (hrR3). The immune model by Al-Tuwairqi et al. (2020) incorporated components of the innate immune response, including natural killer cells and cytokine activity.

**Table 1:** Initial conditions for model 1 variables.

| Variable | Value | Unit | Description |
| --- | --- | --- | --- |
| $C_0$ | $10^6$ | cells/mm<sup>3</sup> | Initial density of uninfected cells |
| $I_0$ | 0 | cells/mm<sup>3</sup> | Initial density of infected cells |
| $V_0$ | $1.9 \times 10^{10}$ | viruses/mm<sup>3</sup> | Initial density of virus particles |
| $Y_0$ | $10^4$ | units/mm<sup>3</sup> | Initial density of immune cells |

**Table 2:** Baseline parameter values for the base model (1) and their corresponding references.

| Parameter | Description | Baseline Value | Units | Source |
| --- | --- | --- | --- | --- |
| $D_C$ | Diffusion coefficient of susceptible cells | 0.006 | $\text{mm}^2$ per day | Baabdulla and Hillen (2024) |
| $D_I$ | Diffusion coefficient of infected cells | 0.006 | $\text{mm}^2$ per day | Baabdulla and Hillen (2024) |
| $D_V$ | Diffusion coefficient of virus | 0.24 | $\text{mm}^2$ per day | Baabdulla and Hillen (2024) |
| $D_Y$ | Diffusion coefficient of immune cells | 0.006 | $\text{mm}^2$ per day | Baabdulla and Hillen (2024) |
| $r$ | Growth rate of uninfected tumor | 0.3 | $(\text{day})^{-1}$ | Baabdulla and Hillen (2024) |
| $L$ | Tumor carrying capacity | $10^6(10^6 - 10^8)$ | cells | Baabdulla and Hillen (2024) |
| $\beta$ | Infection rate of uninfected tumor cells by OV | $1.5 \times 10^{-9}$ | $(\text{PFU} \cdot \text{day})^{-1}$ | Baabdulla and Hillen (2024) |
| $p_1$ | Killing rate of tumor by immune cells | $0.48 \times 10^{-6}$ | $(\text{cell} \cdot \text{day})^{-1}$ | Al-Tuwairqi et al. (2020)(estimated) |
| $\alpha$ | Death rate of infected tumor cells | 1 | $(\text{day})^{-1}$ | Baabdulla and Hillen (2024) |
| $q_1$ | Killing rate of infected tumor by tumor-specific immune | $0.63 \times 10^{-6}$ | $(\text{cell} \cdot \text{day})^{-1}$ | Al-Tuwairqi et al. (2020)(estimated) |
| $b$ | Viruses released by lysed infected tumor cell | 3500 | $(\text{PFU})(\text{cell})^{-1}$ | Baabdulla and Hillen (2024) |
| $\omega$ | Virus-induced immune response rate | 4 | $(\text{day})^{-1}$ | Baabdulla and Hillen (2024) |
| $s_1$ | Killing rate of virus by immune cells | $0.21 \times 10^{-6}$ | $(\text{cell} \cdot \text{day})^{-1}$ | Al-Tuwairqi et al. (2020)(estimated) |
| $\eta_1$ | Stimulation of immune response by uninfected cells | $3.84 \times 10^{-7}$ | $(\text{cell} \cdot \text{day})^{-1}$ | Al-Tuwairqi et al. (2020)(estimated) |
| $\zeta_1$ | Stimulation of immune response by infected cells | $7.8 \times 10^{-7}$ | $(\text{cell} \cdot \text{day})^{-1}$ | Al-Tuwairqi et al. (2020)(estimated) |
| $\xi_1$ | clearance rate of immune cells | 0.036 | $(\text{day})^{-1}$ | Al-Tuwairqi et al. (2020) |

**Table 3:** Parameters of the non-dimensional model 2.

| Parameter | Description | Baseline Value |
| --- | --- | --- |
| $D_c$ | Diffusion coefficient of susceptible cells | 0.025 |
| $D_i$ | Diffusion coefficient of infected cells | 0.025 |
| $D_y$ | Diffusion coefficient of immune cells | 0.025 |
| $\theta$ | Effective viral production rate | 58.33 (16.6 – 500) |
| $p$ | Killing rate of tumor by immune cells | 1.6 |
| $q$ | Killing rate of infected tumor by tumor-specific immune cells | 2.1 |
| $a$ | infected death rate | 3.33 |
| $\gamma$ | Virus clearance rate | 13.33 |
| $s$ | Killing rate of virus by immune cells | 0.71 |
| $\eta$ | Stimulation of immune response by uninfected cells | 1.28 |
| $\zeta$ | Stimulation of immune response by infected cells | 2.6 |
| $\xi$ | clearance rate of immune cells | 0.16 |

## 3 Analysis of the Kinetic Part

We first examine the aforementioned model in the spatially homogeneous scenario, where all spatial dependencies are disregarded. Consequently, the system governing *C*(*t*), *I*(*t*), *V* (*t*), *Y* (*t*) becomes:

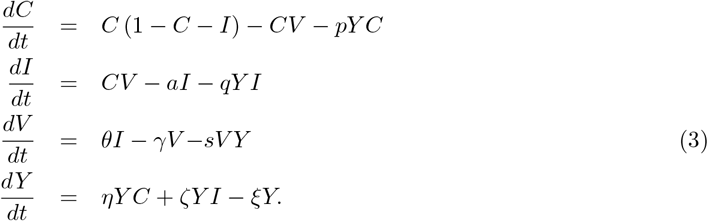

System (3) has five equilibria: The “disease-free” equilibrium is *E*_0_ = (*C, I, V, Y*) = (0, 0, 0, 0), the “cancer-only” equilibrium is given by *E*_1_ = (1, 0, 0, 0), and the “immune-free” equilibrium has the coordinates

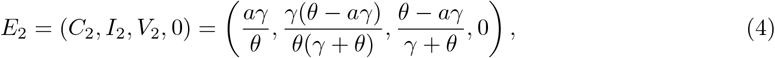

which is only biologically relevant if and only if *θ > γλ*. We denote the “immune-cancer-only” equilibrium as

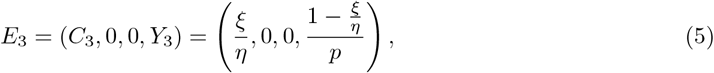

which is biologically relevant if and only if *η > ξ*. The “coexistence” equilibrium is 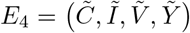 which is a nonzero coexistence point that can not be expressed explicitly due to complexity. To demonstrate the existence of *E*_4_, we will use numerical methods. We simplify the system (2) by assuming that all populations are nonzero. First, we set all four equations to zero. From the fourth equation of system (2), we cancel *Y*, isolate *C*, and obtain

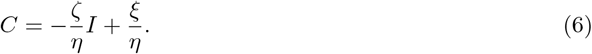

From the second equation of systm (2), isolating *I* gives

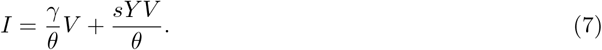

Substituting (7) into (6), we rewrite *C* as

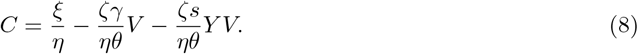

Next, canceling *C* from the first equation of system (2) yields

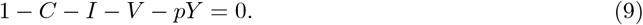

By substituting (7) and (8) into (9), we obtain an equation in terms of two variables, *V* and *Y*:

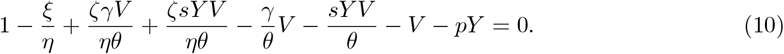

Similarly, substituting (7) and (8) into the third equation of the system (2) gives another relation in *V* and *Y*. Canceling *V* from both sides yields

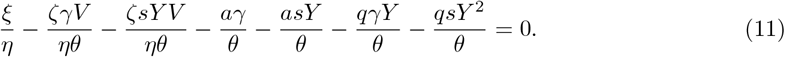

Equations (10) and (11) form a system with two variables. Numerical solutions indicate that this system admits at least one positive solution, establishing the existence of the coexistence point *E*_4_. However, due to the complexity of the equations, explicit expressions cannot be derived, and uniqueness cannot be demonstrated analytically.

The equilibrium points *E*_0_, *E*_1_, and *E*_2_ correspond to the three equilibria of the OV model of Baabdulla and Hillen (2024) and their stability turns out to be the same as in the simpler model. In the *E*_2_ state, the tumor persists while the immune population has been driven to extinction. In the equilibrium point *E*_3_, both infected cells and free viruses tend to zero. This is caused by the immune system responding too quickly to the presence of viruses and infected cells; thus, the immune cells destroy the virus and the infected cells but leave tumor cells behind.

A useful and essential measure in any disease modeling is the basic reproduction number *R*_0_ (Roberts, 2007; Heesterbeek, 2002; Heffernan et al., 2005). The basic reproduction number refers to the expected number of secondary infections resulting from a single primary infection. In virotherapy, primary and secondary infections refer to individual tumor cells infected by the oncovirus. Using the framework of van den Driessche and Watmough (2002), we compute the reproduction number as spectral radius of the next generation matrix *FV*^1^ (Bianchi et al., 2019). The operators *F* and *V* describe new infections and transitions between infected compartments, respectively. The matrices *F* and *V* are the corresponding linearizations at the disease free equilibrium *E*_1_. From model (3) have:

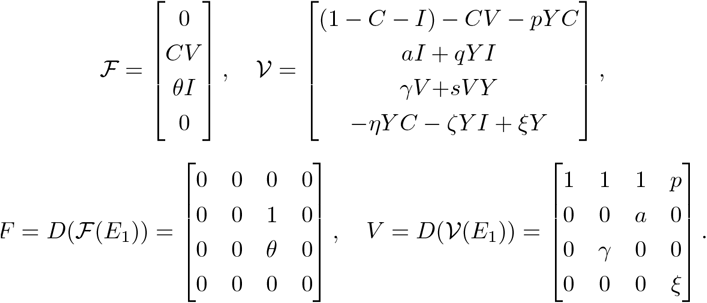

We evaluate the matrix *FV*^1^ and its spectral radius gives us the basic reproduciton number *R*_0_

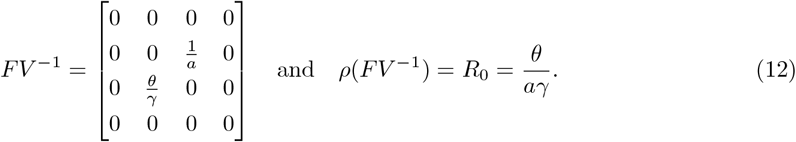

This shows that *E*_1_ is only stable when *R*_0_ *<* 1 which is equivalent to when *θ < aγ*.

We summarize the stability in the following result.

### Theorem 1.

*For our base immune-oncolytic virus model (2), we have the following results:*

- *The basic reproduction number R*_0_ *is given by* 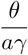,
- *E*_0_ *is always a saddle, and when the basic reproduction number R*_0_ *is less than one or equivalently θ < aγ, E*_1_ *is locally asymptotically stable and E*_2_ *is not biologically relevant*,
- *When R*_0_ *is greater than one, E*_1_ *becomes unstable and the coexistence steady state arises through a transcritical bifurcation at θ*_*t*_ = *aγ, where E*_2_ *becomes biologically relevant*,
- *E*_2_ *is locally asymptotically stable when*

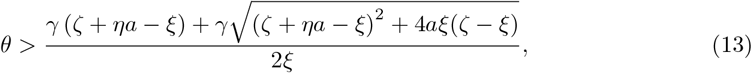

*and*

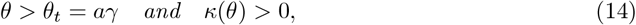

*where*

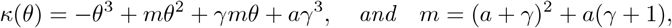
- *There exists a bifurcation value θ*_*H*_ *> θ*_*t*_ *with κ*(*θ*_*H*_) = 0 *such that the system undergoes a Hopf bifurcation at E*_2_,
- *E*_3_ *is locally asymptotically stable only if we have*

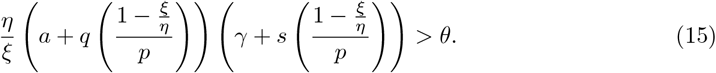

*Proof*. The Jacobian of the system (3) at a general equilibrium 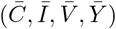 is gisven by

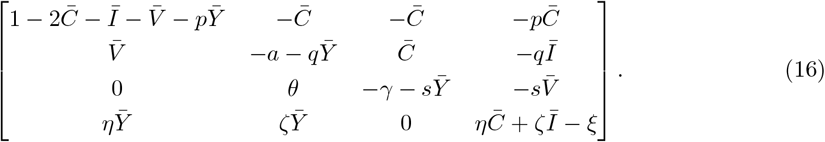

**Part 1**. At *E*_0_ = (0, 0, 0, 0), (16) reduces to

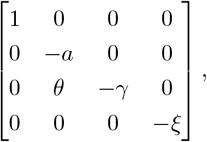

which shows that *E*_0_ is a saddle.

**Part 2**. At *E*_1_ = (1, 0, 0, 0), the Jacobian matrix (16) reduces to

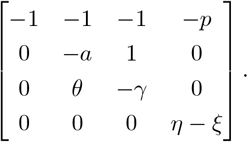

The characteristic equation is given by

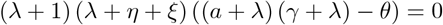

Thus, the eigenvalues are given by *λ*_1_ = −1, *λ*_2_ = *η* − *ξ* and

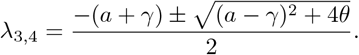

This gives that *E*_1_ is only stable if *ξ > η* and (*γ* + *a*)^2^ > (*γ* − *a*)^2^ + 4*θ*. The second condition is equivalent with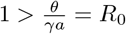.

So far we have shown *E*_0_ is always a saddle and when the condition 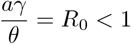 is satisfied, *E*_1_ is locally asymptotically stable and *E*_2_ is not biologically relevant. Howeve*θ*r, when *R*_0_ *>* 1, *E*_1_ becomes unstable and *E*_2_ becomes biologically relevant.

**Part 3**. To analyze the stability of *E*_2_, we set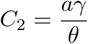, so we rewrite *E*_2_ as

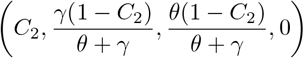

We have that the Jacobian matrix (16) at this point is given by

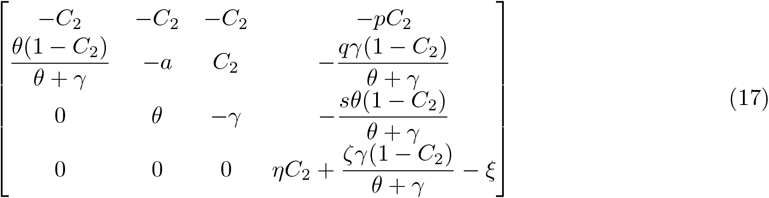

and so the characteristic equation is given by

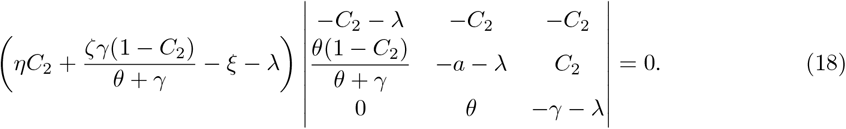

We rewrite this equations as

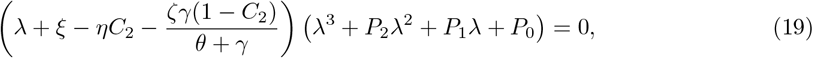

where

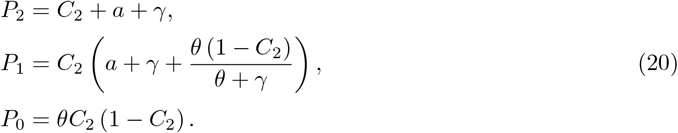

In order to have that *E*_2_ is locally asymptotically stable, we need to show that we have negative real eigenvalues or complex eigenvalues with negative real parts. Let us first disregard the first factor and focus only on the eigenvalues from the second factor (19). We want to analyze the second term to determine the signs of the eigenvalues. We want to show that the polynomial

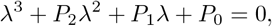

has only real negative eigenvalues, or complex eigenvalues with negative real parts, provided that *θ > aγ* and *κ*(*θ*) *>* 0 where *κ*(*θ*) = − *θ*^3^ + *mθ*^2^ + *γmθ* + *aγ*^3^ and *m* = (*a* + *γ*)^2^ + *a*(*γ* + 1). To see this, we apply the Routh-Hurwitz criterion (Edelstein-Keshet, 2005 - 1988). Based on this criterion for the third order polynomial

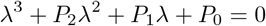

to have that all eigenvalues are either real and negative, or complex with negative real parts, we should have

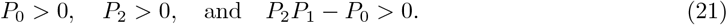

Since from the beginning we had the assumption *C*_2_ *>* 1, we clearly have *P*_2_ *>* 0 and *P*_0_ *>* 0. To check the other condition, we substitute in the values in the *P*_2_*P*_1_ − *P*_0_ and by noting that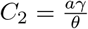, we get

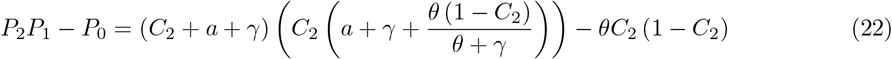

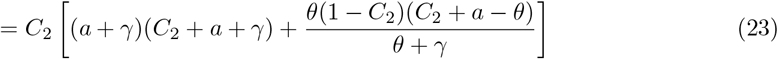

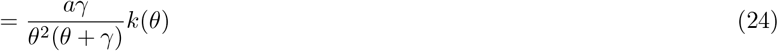

where

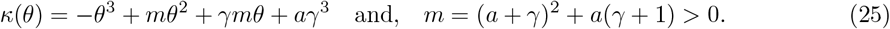

Since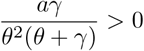, to have the condition *P*_2_*P*_1_ − *P*_0_ *>* 0 satisfied, we should have

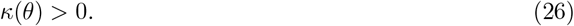

Now we need to analyze the first term in (19) to find all the conditions under which *E*_2_ is locally asymptotically stable. To have this, we need

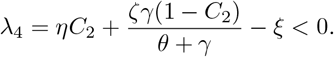

We rewrite this as

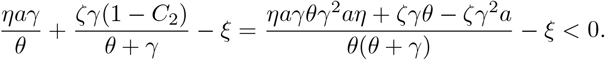

This gives us

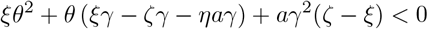

and so to have the condition (26) satisfied, we need

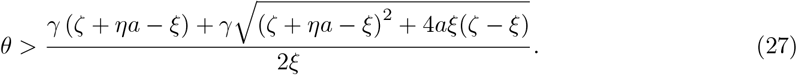

We showed that the condition (21) is satisfied, thus to have *E*_2_ locally asymptotically stable, we need the inequality (27) to be satisfied as well.

**Part 4**. We now want to show that the system (3) undergoes a Hopf bifurcation at *θ*_*H*_ for some *θ*_*H*_ *> θ*_*t*_ for *E*_2_. We again consider the characteristic equation (19). The first factor is always real valued, hence for a Hopf bifurcation, we disregard it and focus on the second part.

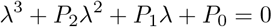

with *P*_2_, *P*_1_, and *P*_0_ values given in (20). We define *H*(*θ*) as

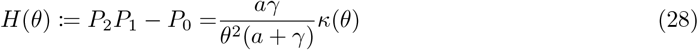

with *κ*(*θ*) defined in (25). We apply Liu’s criterion (Liu, 1994; Jahedi et al., 2021), which states that, assuming *P*_0_ ≠ 0, a Hopf bifurcation occurs at *θ* = *θ*_*H*_ for system (2) whenever

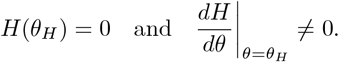

Since the multiplicative factor of *H*(*θ*), *aγ*, is nonzero, it follows that *H*(*θ*) = 0 if and only if *κ*(*θ*) = 0. By Descartes’ rule of signs (Curtiss, 1918), the third order polynomial *κ*(*θ*) has exactly one positive real root, as there is only one sign variation among its coefficients. Considering *κ*(*θ*), there are two sign changes, which implies that *κ*(*θ*) has either zero or two negative real roots. Thus, there exists a unique positive solution *θ* = *θ*_*H*_ such that

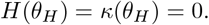

Moreover, since *θ*_*H*_ is the unique positive root of *κ*(*θ*) and the leading-order term of *κ*(*θ*) is negative, we conclude that *κ*(*θ*) *<* 0 for all *θ > θ*_*H*_. Consequently, because *H*(*θ*) in Equation (28) has a positive multiple, *aγ >* 0, we also have *H*(*θ*) *<* 0 for all *θ > θ*_*H*_. Finally, from (26), we know that *κ*(*θ*) *>* 0, which implies *θ*_*t*_ *< θ*_*H*_.

**Part 5**. To analyze the stability of *E*_3_ = (*C*_3_, 0, 0, *Y*_3_), we have the Jacobian matrix given by

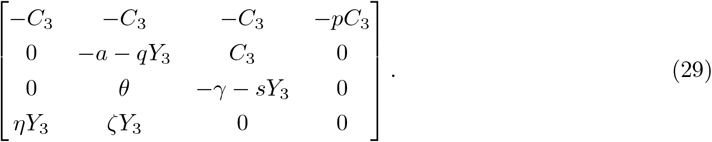

The characteristic equation is given by

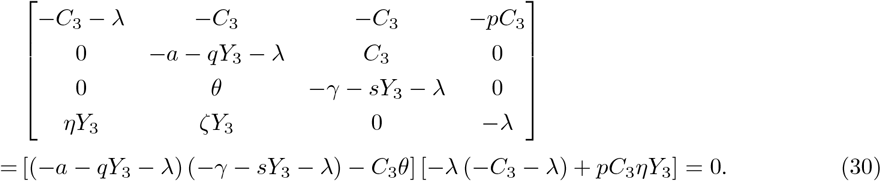

Considering the cancer-immune equilibria *E*_3_ = (*C*_3_, 0, 0, *Y*_3_) (5), we substitute the values of *C*_3_ and *Y*_3_ into the Equation (30). The second factor gives us

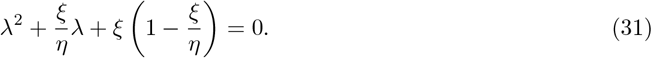

The sum of the roots of this quadratic equation is given by the negative of the coefficient of the linear term, i.e. 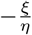, which is strictly negative. The product of the roots of this equation is equal to the constant term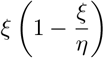,, which is positive since we have *ξ < η*. This means that if this quadratic equation has real roots, they are negative. Thus, the eigenvalues from this factor of the characteristic equation (30) would be negative.

We follow the same approach for the first factor of (30). After substituting the values of *C*_3_ and *Y*_3_ into the first factor of Equation (30), we obtain

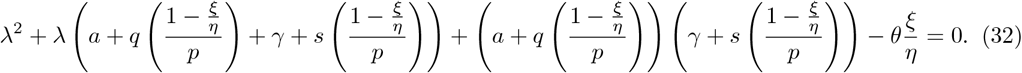

Similarly, the sum of the roots of this equation is equal to the negative of the coefficient of the linear term, i.e.

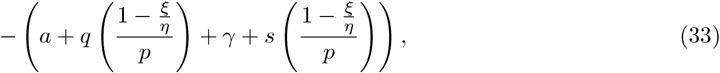

which has a negative value. If real roots exist, then for them to be negative we need to have their product positive. To have this condition satisfied, we require the constant term of the Equation (32) to be positive, that is

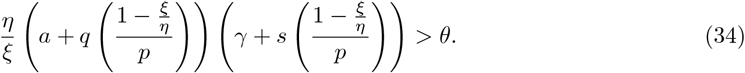

Hence *E*_3_ is locally asymptotically stable only if we have (15).

We also consider *η* as a bifurcation parameter and for low virus concentration we find a transcritical bifurcation in *η*.

### Lemma 2.

*Assume the virus concentration V and the infected population I are close to* 0. *Then by increasing η we obtain a transcritical bifurcation between E*_1_ *and E*_3_ *at η* = *ξ*.

*Proof*. To study the transcritical bifurcation between the equilibria *E*_1_ = (1, 0, 0, 0) and *E*_3_ = (*C*_3_, 0, 0, *Y*_3_), we set *I* = 0 and *V* = 0. In this case, system (2) reduces to

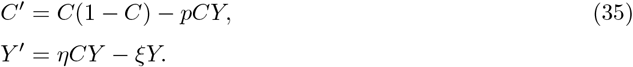

The steady states of this reduced system are: the trivial equilibrium *e*_0_ = (0, 0), the cancer-only equilibrium *e*_1_ = (1, 0), and the coexistence equilibrium 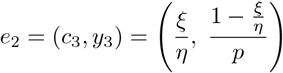. These equilibria are consistent with those found in the full system (2) when the infected population *I* and the virus population *V* are set to zero.

The Jacobian of system (35) at a steady state 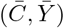 is

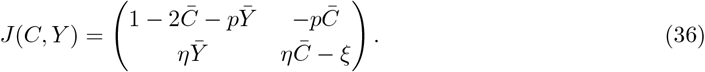

At the trivial equilibrium (0, 0), the system is always a saddle. At the cancer-only equilibrium (1, 0), the Jacobian is

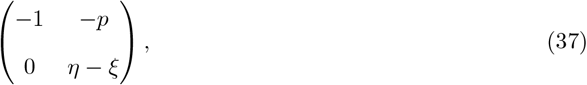

which shows that (1, 0) is stable when *ξ > η*. The condition *ξ* = *η*, therefore, defines a bifurcation point.

At the coexistence equilibrium *e*_2_ = (*c*_3_, *y*_3_), the Jacobian becomes

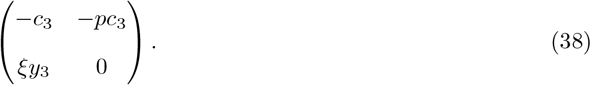

The eigenvalues are given by

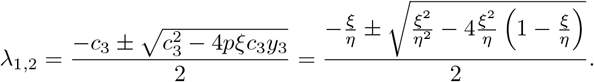

For *ξ < η*, the real parts of the eigenvalues are negative, while for *ξ > η*, the equilibrium *e*_2_ becomes a saddle. Hence, a transcritical bifurcation occurs between *e*_1_ and *e*_2_ at *ξ* = *η*.

In Figure 1, we plot the curve

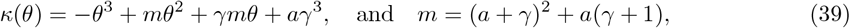

using the parameters values from Table 3. We have indicated the points *θ*_*t*_ = *aγ* and the Hopf bifurcation value *θ*_*H*_ on this curve. Their corresponding values are 44.4 and 338.4, respectively.

**Figure 1:**
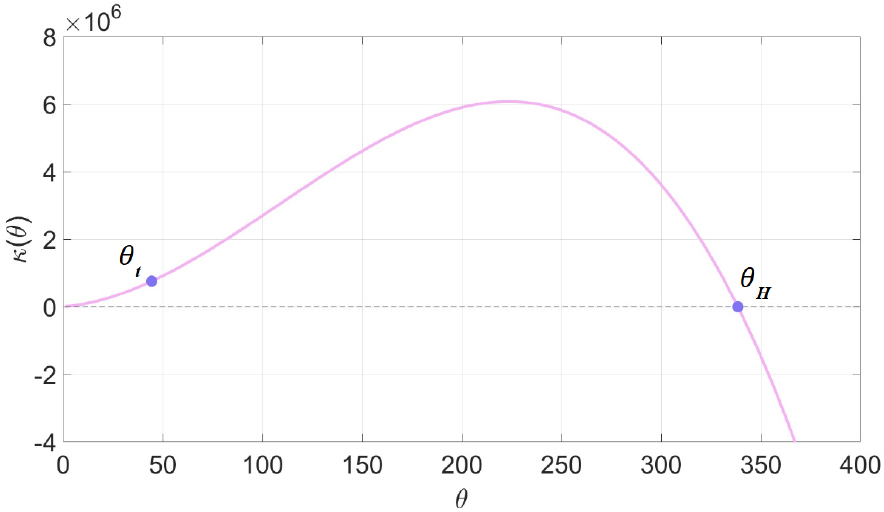
The *κ*(*θ*) curve of Equation (25), with the parameter values from Table 3 the bifurcation values *θ*_*t*_ = 44.4 and *θ*_*H*_ = 338.45 are indicated on the curve.

In Figure 2, we illustrate the stability behavior of system 2 for various values of *θ* and *η*, with *θ* ranging from 30 to 500 and *η* from 0 to 2. Each colored dot represents the stability type of a steady state within the given parameter region. A red dot indicates that the cancer-only equilibrium *E*_1_ is stable for the corresponding range of parameter values. Green dots mark the regions where *E*_3_ is stable, while light blue denotes the stability of *E*_2_. For *θ* values greater than the Hopf bifurcation threshold *θ*_*H*_, we have the stable limit cycles around *E*_2_, shown in dark blue. Yellow dots represent the stability of *E*_4_, and for certain parameter values, we observed numerically stable limit cycles around *E*_4_, which are illustrated in orange.

**Figure 2:**
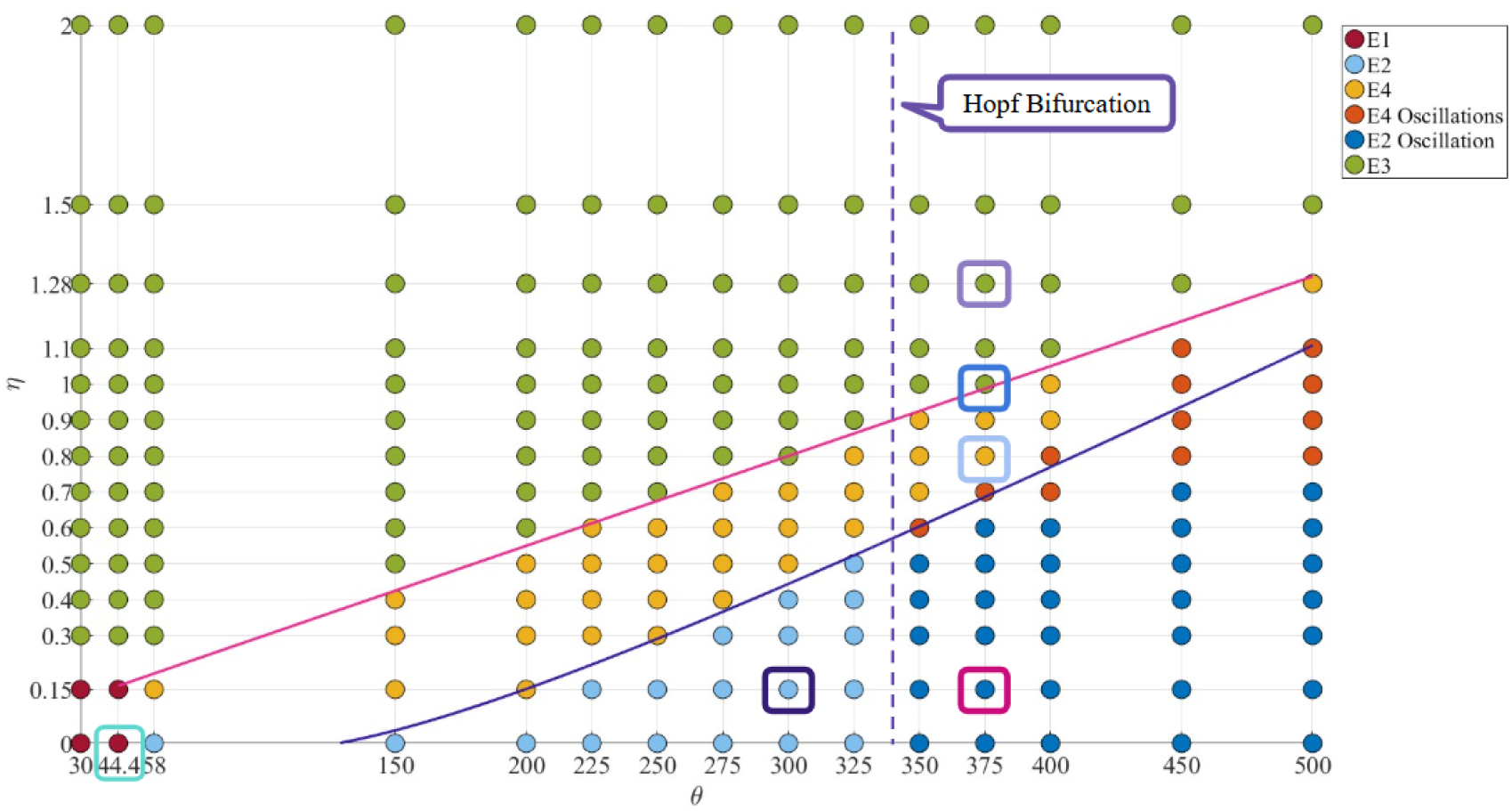
Stability behavior of system (1) over a range of *θ* and *η* values. The dashed line indicates the bifurcation value *θ*_*H*_ for the equilibrium *E*_2_. The equations given in (40) and (41) are represented by the pink and blue curves. The squared markers highlight parameter values corresponding to the dynamics shown in Figures 3 and 4.

Using inequality (34), we plot the curve *θ*_1_, given by

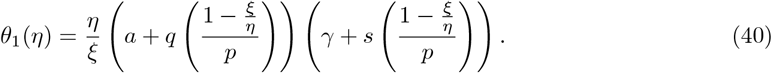

For all *η* values above the *θ*_1_ curve (shown as a pink line), the equilibrium *E*_3_ is stable up to the Hopf Bifurcation value *θ*_*H*_. These points are indicated by green dots. Above the line *θ*_1_(*η*) the virus is cleared and we end up with a cancer-immune equilibrium. Hence to have any hope of successful viro therapy, we need to be below the line *θ*_1_(*η*). Similarly, from inequality (27), we plot the curve *θ*_2_, defined as

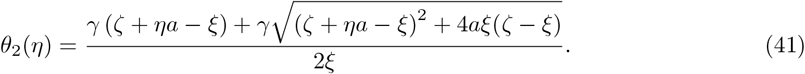

Below the *θ*_2_ curve (shown as a purple line), for *θ* between 30 and *θ*_*H*_ = 338.45, *E*_2_ is stable, indicated by light blue dots. After the Hopf bifurcation values, shown as a dashed line, the point under the *θ*_2_ line numerically show a stable limit cycle around *E*_2_, marked by darker blue dots. In the region between the *θ*_1_ and *θ*_2_ curves, the system exhibits stability of the equilibrium *E*_4_, represented by yellow dots. For higher *θ* values within this region, oscillatory dynamics appear around *E*_4_; the exact transition threshold can not be determined explicitly, and these points are shown in orange.

We select a representative point from each stability region and simulate the system dynamics for these cases, as shown in Figures 3 and 4. The selected points are marked with squares in Figure 2 and are plotted in Figures 3 and 4 using their corresponding stability colors. Figures 3 and 4 present three-dimensional dynamics of system 2 with the variables *C, V*, and *Y* on the axes. We do not show *I* as it closely follows the *V* dynamics. In Figure 3(a), we show the dynamics of system 2 for *η* = 0 and *θ* = 44.4. For these parameter values, Figure 2 indicates that the cancer-only equilibrium *E*_1_ is stable, and hence the system converges to (1, 0, 0, 0). The corresponding simulation of population dynamics is shown in Figure 3(b). In Figure 3(c), we observe the dynamics of system 2 for *η* = 0.15 and *θ* = 300. For these parameter values, from what we see in Figure 2, we have that the immune-free equilibrium *E*_2_ is stable, and the system converges to this point. Its corresponding population dynamics simulation is shown in Figure 3(d). Keeping *η* = 0.15, after *θ* passes the Hopf bifurcation value *θ*_*H*_, for *θ* = 400 we see in Figure 3(e) a stable limit cycle around *E*_2_. We see the corresponding oscillations in Figure 3(f). The system converges to the cancer-immune-only equilibria for *η* = 1.28 and *θ* = 400, as we observe in Figure 4(a), and the population dynamics simulations in Figure 4 (b). Although we can not have the coexistance equilibrium point *E*_4_ explicitly, from the simulations we observe that the system converges to a non-zero equilibrium point for *η* = 1 and *θ* = 400, if both the three-dimensional and population dynamics simulations in Figures 4(c) and 4(d). A stable limit cycle around *E*_4_ is observed for *η* = 0.7 and *θ* = 400 in Figure 4(e) and the oscillatory behavior in Figure 4 (d).

**Figure 3:**
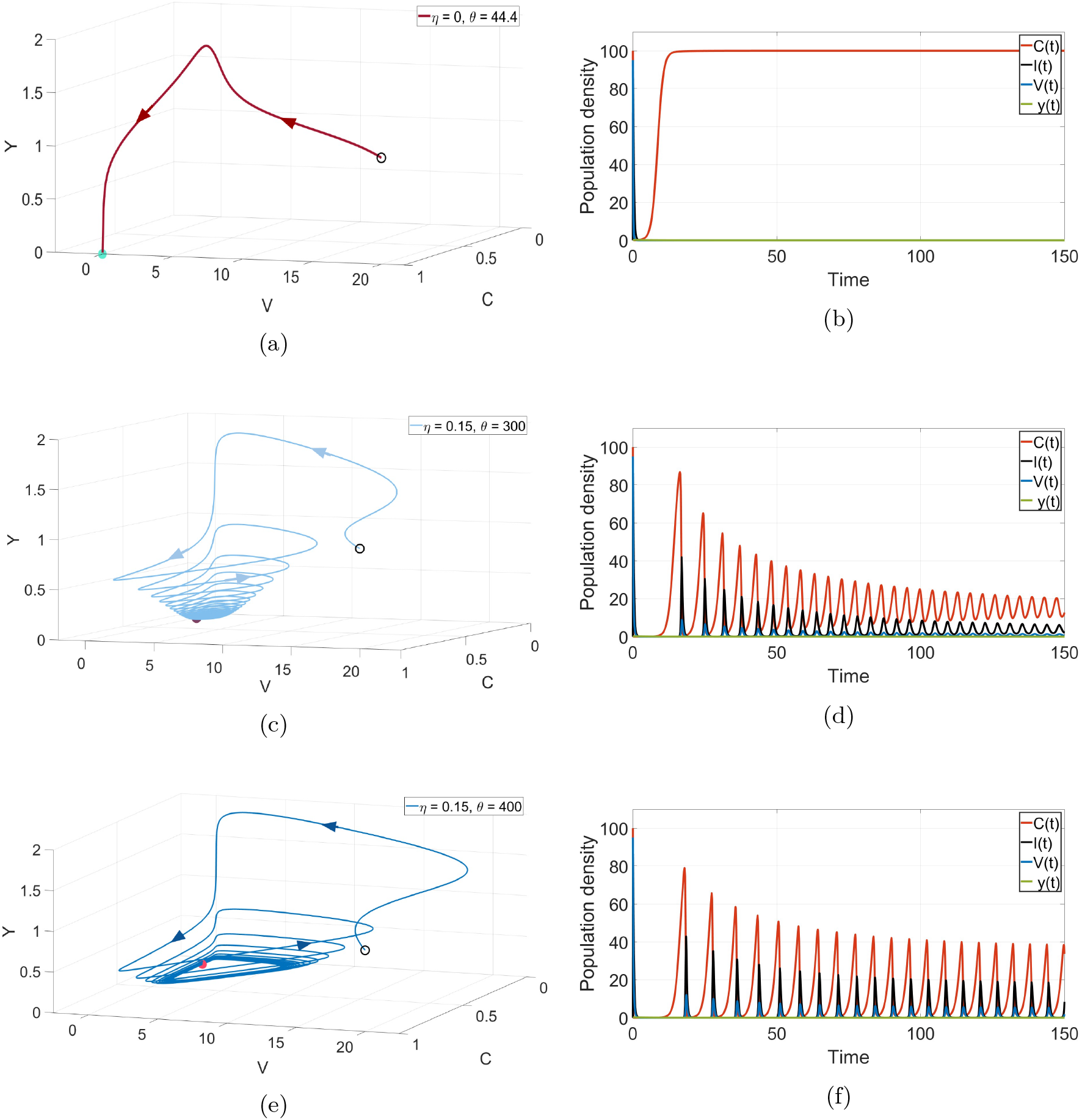
Dynamics of the system (3) for various values of *θ* and *η*. The values used for each simulation are shown in the legends of the left-hand plots. The corresponding two-dimensional time dynamics, with parameter values, are indicated on the right. (a) and (b): the equilibrium *E*_1_ is stable. (c) and (d): the equilibrium *E*_2_ is stable for *θ < θ*_*H*_, prior to the Hopf bifurcation. (d) and (e): A stable limit cycle emerges around *E*_2_ for *θ > θ*_*H*_.

**Figure 4:**
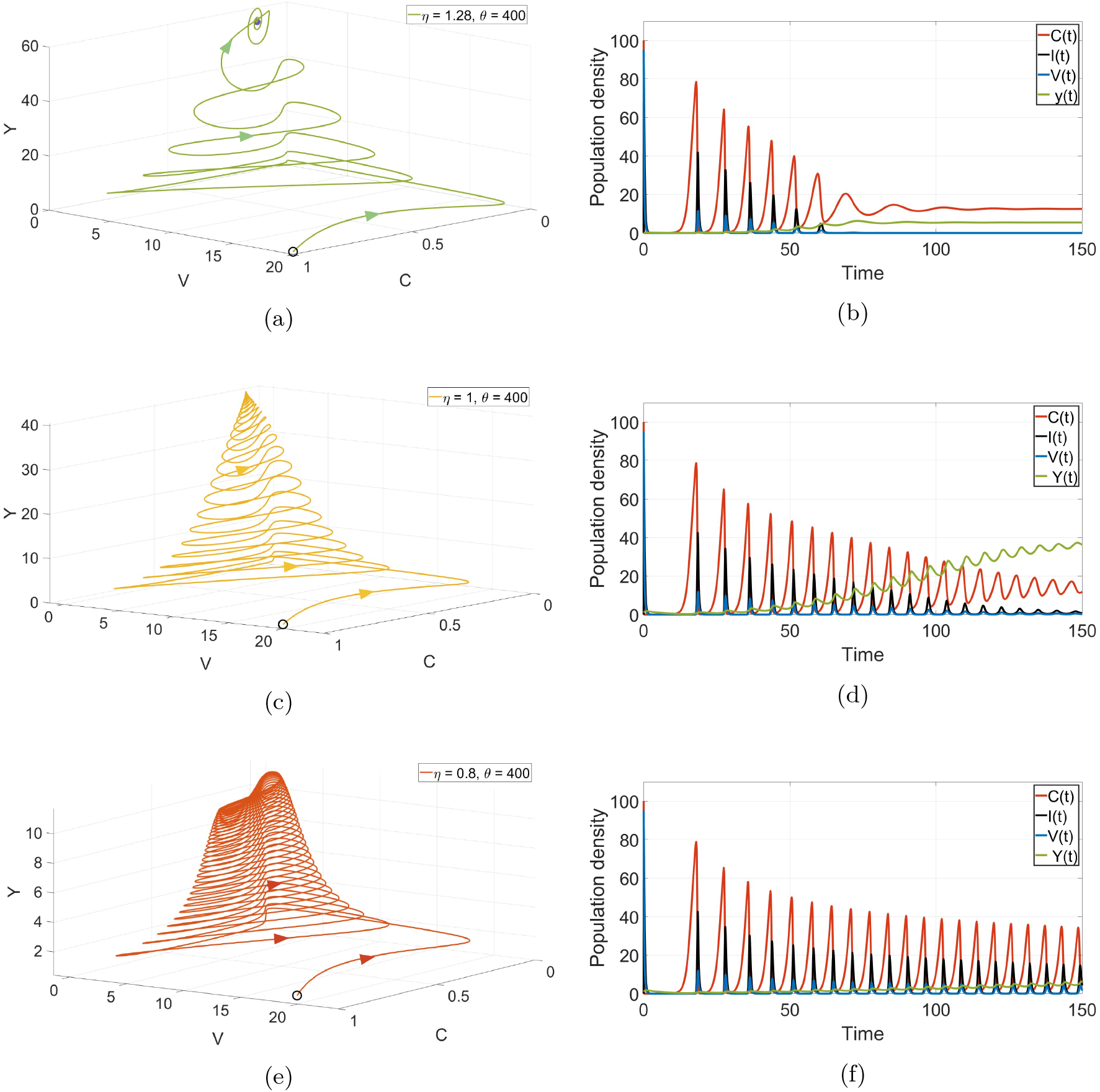
Dynamics of the system (3) for various values of *θ* and *η*. The values used for each simulation are shown in the legends of the left-hand plots. The corresponding two-dimensional time dynamics, with parameter values, are indicated on the right. (a) and (b): the equilibrium *E*_3_ is stable. (c) and (d): the equilibrium *E*_3_ is stable, prior to the Hopf bifurcation. (d) and (e): A stable limit cycle emerges around *E*_4_.

### 3.1 Sensitivity Analysis

We perform a parameter sensitivity analysis with the base value parameters from Table 3. We use this analysis to identify the parameters that most significantly contribute to treatment efficacy. The relative sensitivity of variable *C* to the parameter *p* is defined as (Ingalls, 2013)

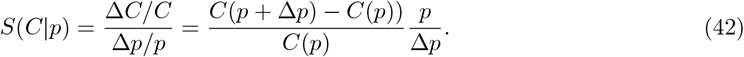

The relative sensitivity relates the size of a relative perturbation in *p* to a relative change in *C*. We vary each parameter by 5% of its value, and we obtain the sensitivity graph as in Figure 5. The sensitivity analysis highlights the crucial role of the immune system in determining the success of oncolytic virotherapy (OVT). While the parameter *θ* has a relatively strong influence compared to most other parameters, its effect is less pronounced than that of *η* and *ξ*. The effectiveness of OVT depends significantly on these two parameters. This suggests that therapeutic strategies should aim to modify parameters of highest sensitivity that most strongly improve treatment outcomes, i.e. the immune stimulation *η*, immune exhaustion *ξ* and the effective production rate of the virus, *θ*.

**Figure 5:**
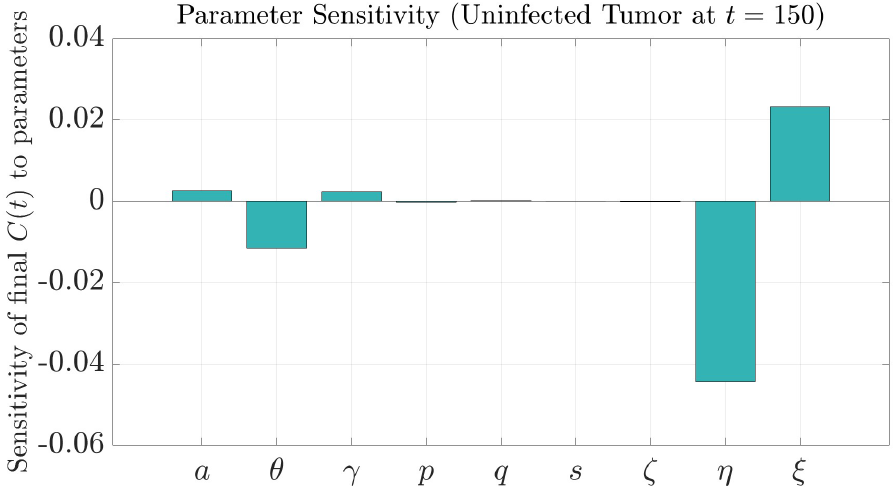
Sensitivity analysis of the final value of *C*(*t*) at *t* = 150 under a 5% change in the model parameter values.

They can be modified through strategies such as combining OVT with immunotherapies. As future work, we propose to investigate which combination strategies are most effective in enhancing treatment outcomes.

## 4 Travelling Fronts and Stacked Waves

In this section, we come back to the spatial model (2) and analyse the invasion speeds of the various components. The model in one dimension reads

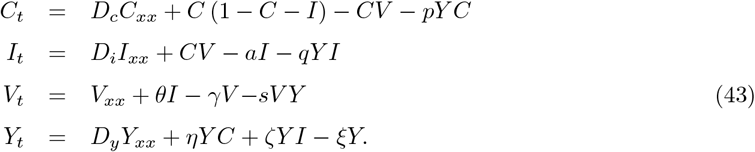

We first simulate the travelling waves in Matlab to observe their behaviour over time. We choose parameter values from Table 3 and diffusion coefficients as *D*_*c*_ = *D*_*i*_ = *D*_*y*_ = 0.025. The initial conditions for *I* and *V* are set at *E*_2_, with *Y* chosen to be equal to a small value, 0.18, for *x* between 0 and 4 on the left boundary, and zero elsewhere. The uninfected cancer population *C* is initialized at its carrying capacity up to *x* = 30, and 0 elsewhere, representing the invasion of cancer into the spatial domain. This setup allows us to illustrate the invasion of cancer into the healthy domain *x >* 30. As shown in Figure 6, the leading edge of the wave represents the invasion of uninfected cancer cells (*C*) into healthy tissue. This is followed—at a higher speed—by the virus and infected cells overtaking the uninfected population, reducing the cancer population, until it reaches the wave front, driving the system toward the steady state *E*_2_, where the immune population is at zero. Further back, at a much slower speed, the immune system begins invading the other three populations. Over time, it reduces both the virus and infected cell populations to zero, ultimately settling into a coexistence state with a nonzero cancer population, corresponding to the steady state *E*_3_. We performed many more simulations (not shown here) and this behavior is quite robust. We clearly identify a sequence of invasion fronts that all move at different speeds, i.e. stacked waves. We analyse these stacked waves by looking at the linear invasion speeds of the system for each of the equilibria in turn.

**Figure 6:**
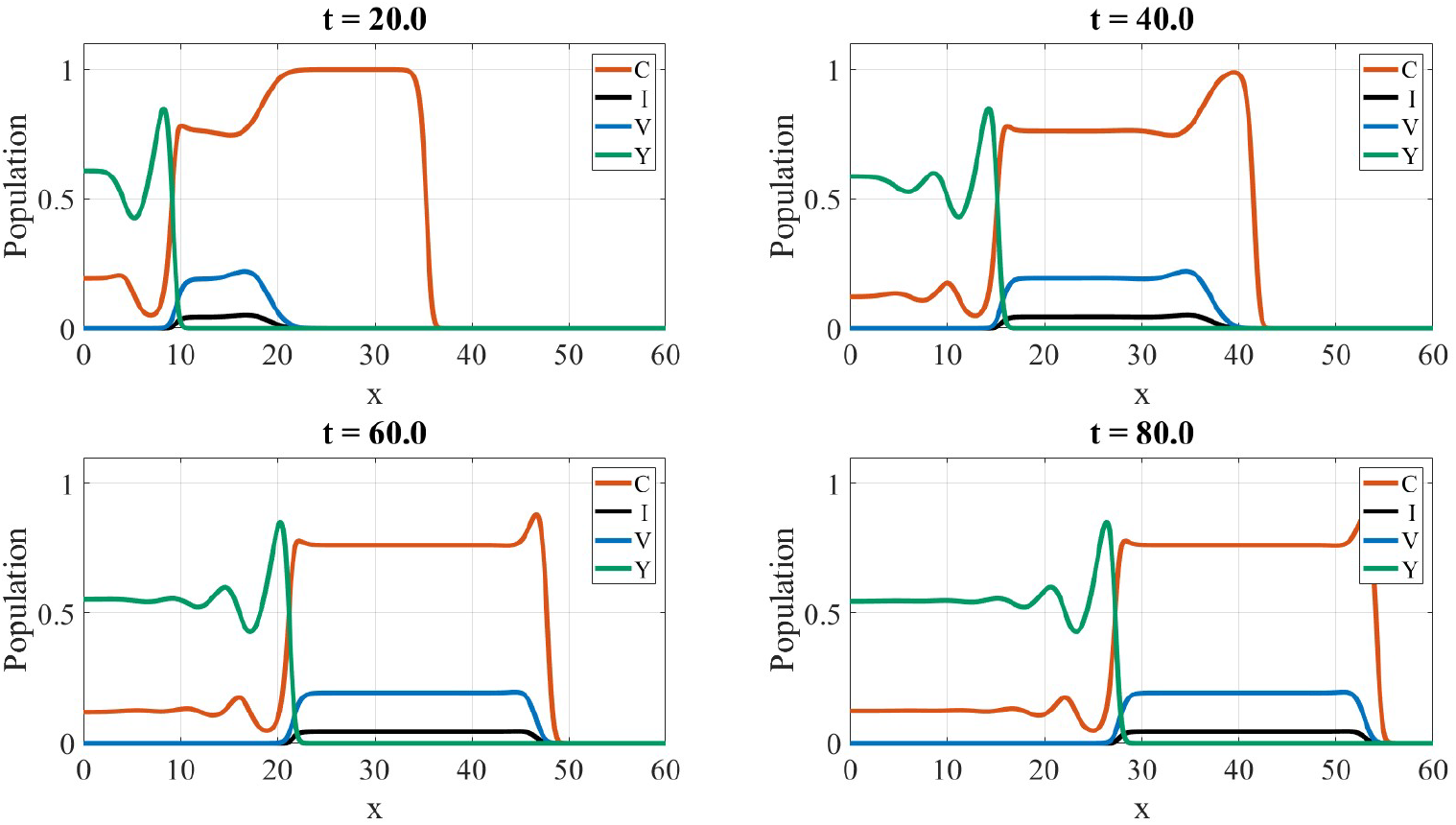
The traveling waves of the four populations invading to the right. The waves are captured at *t* = 20, 40, 60, and 80.

### 4.1 Invading the Disease-Free Equilibrium *E*_0_

To compute the invasion wave front of the cancer into the disease-free equilibrium, we simply consider the cancer-only model

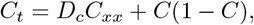

with boundary condition

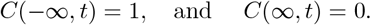

This is the famous Fisher-KPP equation (Vries et al., 2006) and the invasion speed is well known and given by

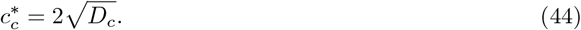

Using the parameter values from Table 3, Equation (45) gives us the minimum speed value

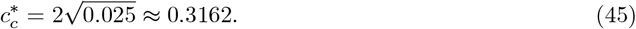

We show this invasion front in Figure 8a.

### 4.2 Invading the Cancer-Only Equilibrium *E*_1_

To consider the invasion of the virus population into an established tumor, we linearize the system in one dimension at the homogeneous steady state (1, 0, 0, 0)

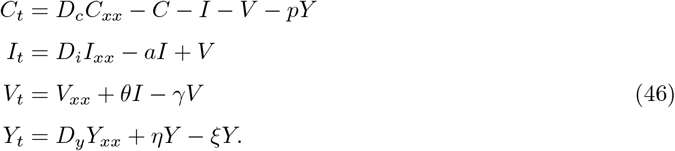

We now make an explicit ansatz of an exponentially decaying self-similar wave solution by setting *z* = *x* − *ct* as

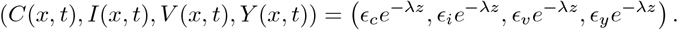

Here, *c* denotes the wave speed, and *λ* the exponential decay rate at the wave front. Substituting this ansatz into system (46), we get

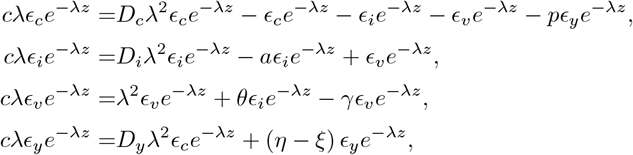

which, after simplification, is written as a linear system

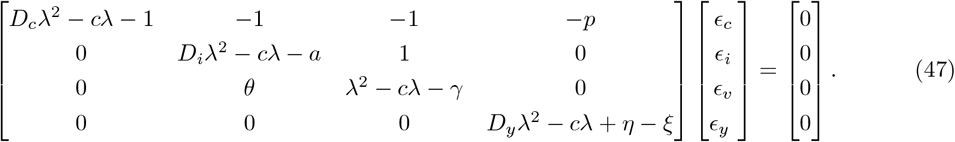

The characteristic equation of the matrix (47) is

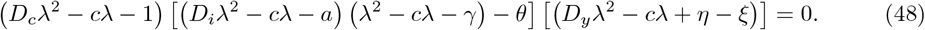

For the first factor of Equation (48), we set

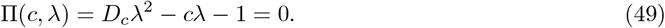

By calculation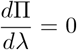, isolating for *c*^′^, and setting it equal to zero, we obtain the minimum value for *c*. Plugging this value back to the Equation (49) yields *D λ*^2^ 1 = 0. For no value of *λ*, the equation can be satisfied and it shows that this factor does not give us any suitable minimum wave speed. We now focus on the second factor of Equation (48). We let

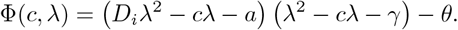

We set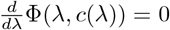 which is given by

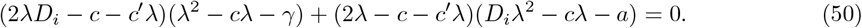

We solve for *c*^′^ and we obtain

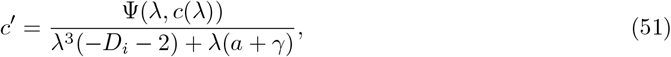

where

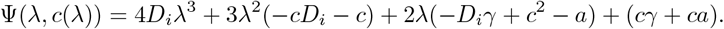

The minimum wave speed 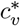 is the value where Φ = 0 and Ψ = 0. We show the functions Φ(*θ*) and Ψ(*θ*) in Figure 7. Using the parameter values from Table 3, we numerically solve for the minimum wave speed as

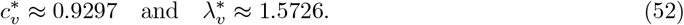

**Figure 7:**
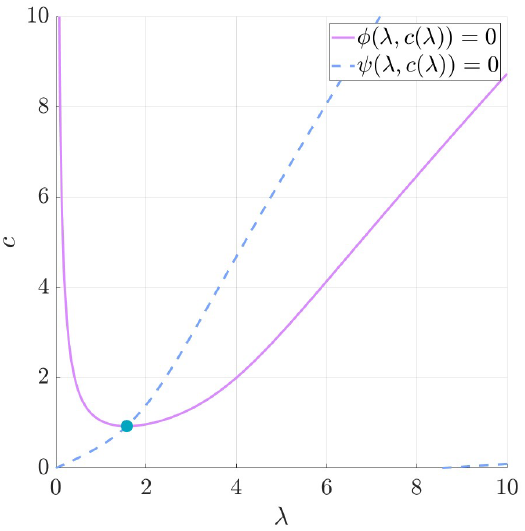
The blue ball corresponds to the minimum decay rate and minimum wave speed of the virus particles.

**Figure 8:**
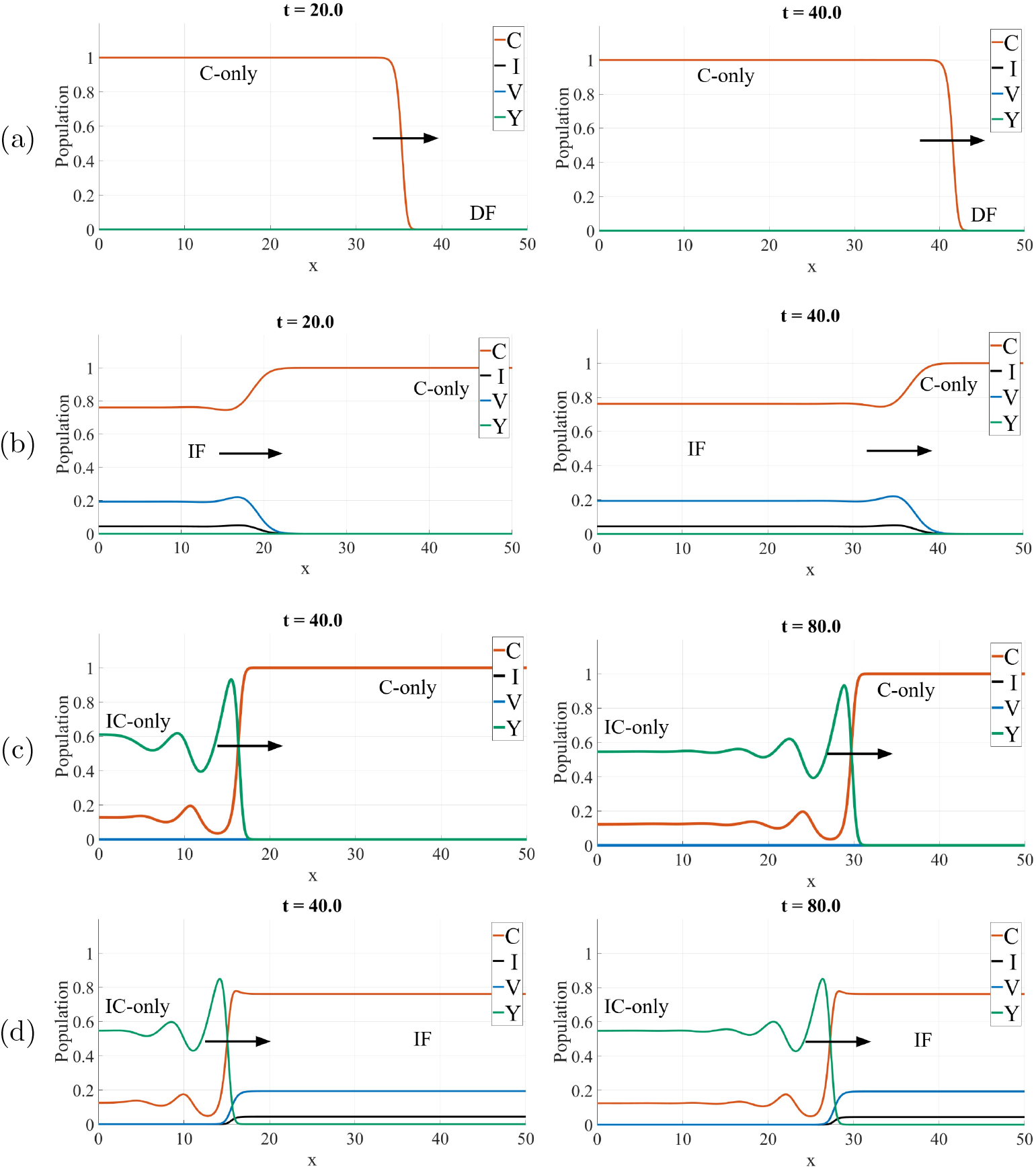
Traveling waves of the different populations at various times: (a) The cancer-only (C-only) wave invading to the right to the disease-free (DF) equilibria. The wave moved from *X*_1_ = 35.298 in *t* = 20 to *X*_2_ = 41.541 at *t* = 20. (b) The imune-free (IF) equilibria invading the Cancer only (C-only). The wave moved from *X*_1_ = 18.969 in *t* = 20 to *X*_2_ = 36.978 at *t* = 20. (c) The immune-cancer only (IC-only) invading cancer-only(c-only) equilibria. The wave moved from *X*_1_ = 16.28 in *t* = 40 to *X*_2_ = 29.695 at *t* = 80. (d) The immune-cancer only (IC-only) invading immune-free(IF) equilibria. The wave moved from *X*_1_ = 15.048 in *t* = 40 to *X*_2_ = 27.254 at *t* = 80.

This speed corresponds to the invasion of virus and infected cancer cell populations into an established cancer and is illustrated in Figure 8b. There is a third term in equation (48), which we analyse next.

We use the same approach by defining *ϕ*(*λ, c*) as

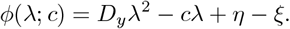

By setting *ϕ*(*λ*; *c*) = 0, we obtain

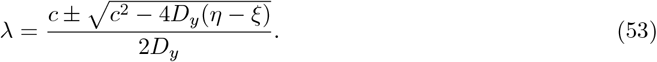

We find the minimum wave speed values *c*(*λ*), as the minimum value of *c* for which *λ* is real. Thus, we have

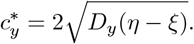

Using the parameters from Table (3), this value corresponds to:

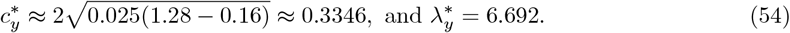

This speed describes the invasion of immune cells into an established tumor. A simulation is shown in Figure 8c.

### 4.3 Invading the Immune-Free Equilibrium *E*_2_

This case corresponds to the speed with which the immune system invades the immune-free equilibrium state *E*_2_. We linearize (17) at *E*_2_ = (*C*_2_, *I*_2_, *V*_2_, 0).

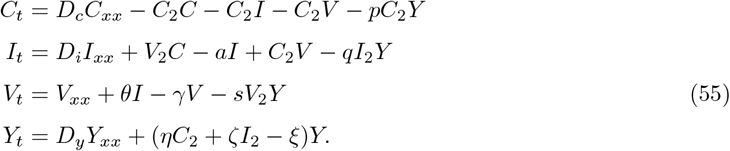

Similarly to the previous case, we make an explicit ansatz for an exponentially decaying self-similar wave solution by setting *z* = *x* − *ct* as

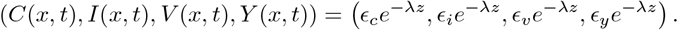

Substituting this ansatz into system (55) and writing it as a linear system we obtain

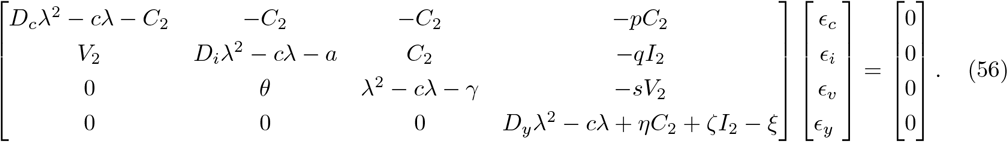

The characteristic equatin is given by

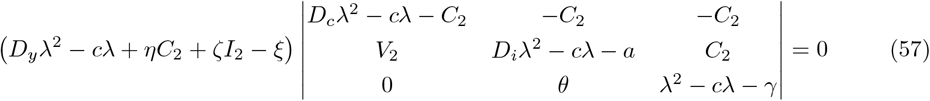

Let us define *χ*(*λ, c*) as the first factor

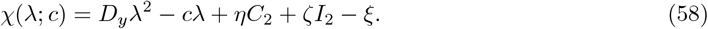

By setting *χ*(*λ*; *c*) = 0, we obtain

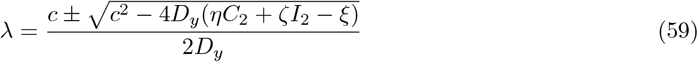

We find the minimum wave speed values *c*(*λ*), as the minimum value *χ* for which it has a real solution. Thus, we have

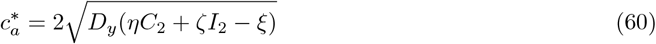

Using the parameters from Table 3, we have this value corresponding to:

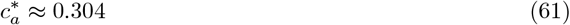

### 4.4 Numerical Simulations

To obtain the travelling speeds of the four waves numerically and compare them with the theoretical values derived above, we set up each wave in a separate simulation. In Figure 8a, the uninfected cancer population is set at its carrying capacity on the left, while all other populations are set to zero. This allows us to calculate the invasion speed of the cancer front moving to the right into healthy tissue. The arrow is used as a marker to track the wave’s position. By measuring the displacement of the marker from *t* = 20 to *t* = 40, we find Δ*X* = 6.20, corresponding to a wave speed of 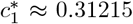,which agrees closely with the theoretical value *c*^*^ 0.3162 in (45). For the second wave, corresponding to the virus invasion speed into the uninfected cancer population (Fig. 8b), we set the right boundary to the steady state *E*_1_ = (1, 0, 0, 0), while the left boundary is set to the immune-free equilibrium *E*_2_. The arrows indicate how the virus front advances into the *C* population, reducing its value and driving the system toward the steady state *E*_2_, where the immune population is absent. From *t* = 20 to *t* = 40, the wave advances by Δ*X* = 18.009, giving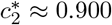, which is in good agreement with the theoretical estimate 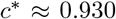 (52). For the third wave (Fig. 8c), we simulate the immune system invading the cancer-only equilibrium *E*_1_. Similarly, the arrows show the immune front, which reduces the *C* to a nonzero value, consistent with the no-virus steady state *E*_3_. Between *t* = 40 and *t* = 80, the marker advances Δ*X* = 13.415, giving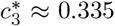, which matches well with the theoretical value in 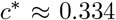 (54). In Figure 8d, we simulate the immune system invading the other populations in immune-free equilibrium *E*_2_. Similarly the arrows show the immune front, which eliminates the *V* and *I* populations and reduces *C* to a nonzero value, consistent with the steady state *E*_3_. Between *t* = 40 and *t* = 80, the marker advances by Δ*X* = 12.206, giving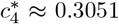, which matches well with the theoretical value in *c*^*^ 0.304 (54).

A simulation of the full stacked wave for this example is shown in the earlier Figure 6. Here the virus infection is the fastest wave and it catches up with the growing tumor after some time. The immune response is rather slow, even slower than the cancer invasion front (see Table 4), hence, in this example, the immune response would have a minimal effect on the virus infection of the cancer. This case contradicts earlier studies, where it was suggested that the immune response removes the virus before it infects all cancer cells.

**Table 4:** Comparison of theoretical and numerical wave speeds for slow versus fast immune response.

| Case | Wave | Theoretical | Numerical |
| --- | --- | --- | --- |
| Slow immune | cancer invasion | $c_c^* = 0.316$ | $c_{c_n}^* = 0.312$ |
| | virus invasion into the cancer | $c_v^* = 0.930$ | $c_{v_n}^* = 0.900$ |
| | immune invasion into cancer | $c_y^* = 0.335$ | $c_{y_n}^* = 0.335$ |
| | immune invasion into virus | $c_a^* = 0.304$ | $c_{a_n}^* = 0.304$ |
| Fast immune | cancer invasion | $c_c^* = 0.316$ | $c_{c_n}^* = 0.312$ |
| | virus invasion into the cancer | $c_v^* = 0.930$ | $c_{v_n}^* = 0.900$ |
| | immune invasion into cancer | $c_y^* = 0.335$ | $c_{y_n}^* = 0.335$ |
| | immune invasion into virus | $c_a^* = 1.058$ | $c_{a_n}^* = 1.032$ |

However, changing model parameters, we make the immune response to invade faster. For example, it would be reasonable to assume that the immune cells move much quicker than the cancer cells. So far we used the same diffusion coefficients for immune and cancer cells (see Table 3). However, if we increase the diffusion coefficient of the immune system by 10 times, we achieve the speed for immune invasion (54) as

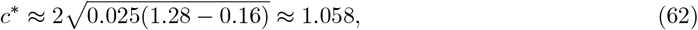

which is faster than the virus invasion speed.

We show simulations of this case in Figure 9, we observe that the immune system quickly catches up with the viral infection, removes the virus and reaches the end of the cancer region. Estimating the immune invasion speed numerically we observe that the immune system advances Δ*X* = 20.658 from *t* = 15 to *t* = 35 which gives the speed of *c*^*^ 1.032 which is a great estimate of the theoretical speed we obtained in (62) *c*_*_ 1.058. Since we only made the change in the diffusion coefficient of the immune cells, the speed of invasion for the other populations stay the same.

**Figure 9:**
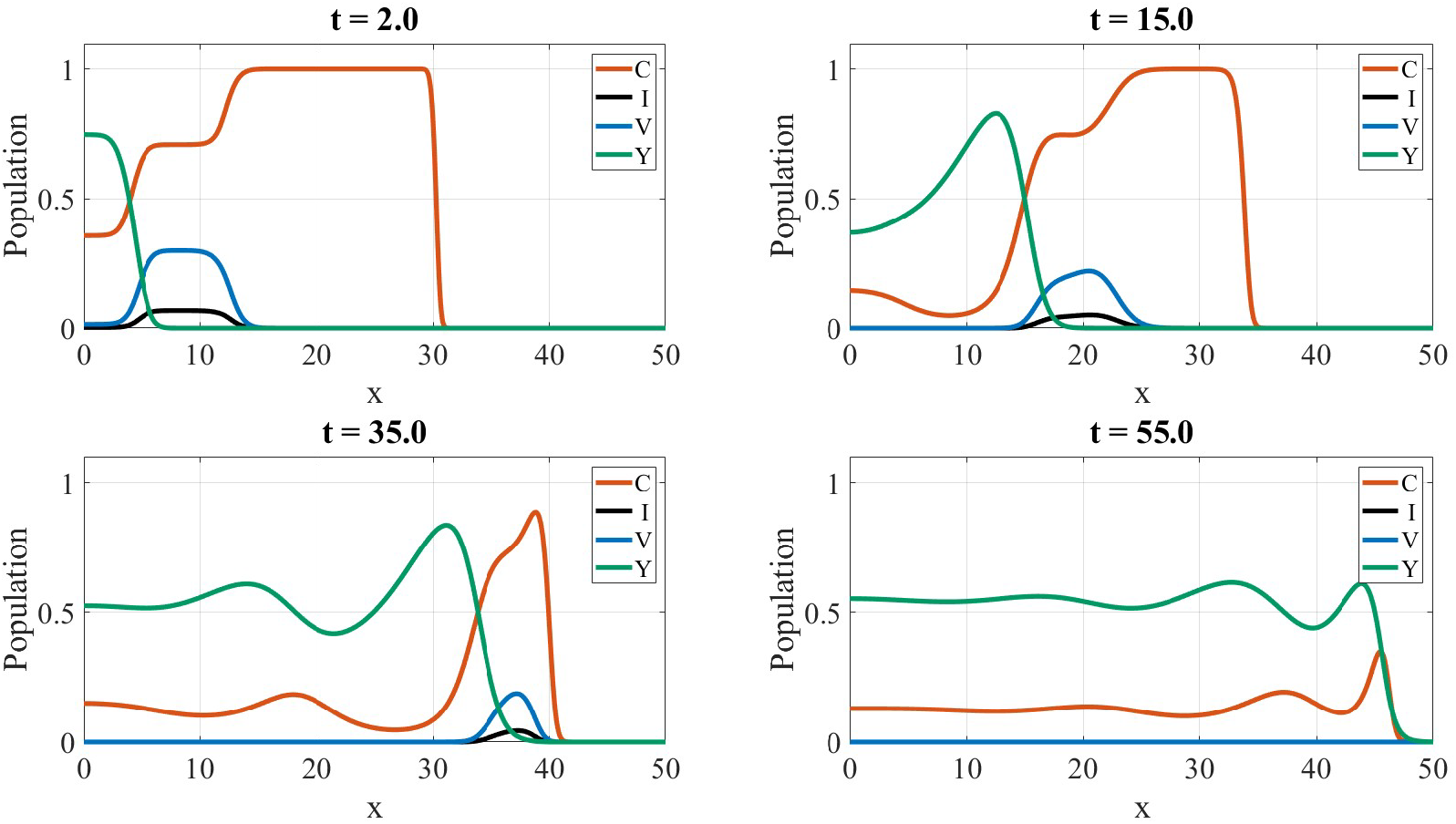
The traveling waves of the four populations invading to the right. The waves are captured at *t* = 2, 15, 35, and 55.

## 5 Conclusion

Using a virus as a cancer treatment is a intriguing idea. In fact, some of these viruses have already passed clinical trials and have obtained FDA approvals for treatment (Greig, 2016). However, oncolytic virotherpay has not reached its maximum efficacy (Russell and Barber, 2018) due to physical barriers, tumor heterogeneity, and an immunosuppressive tumour microenvironment. To further investigate the role of the immune response for OV, we developed a reaction-diffusion model, inspired by the models by Baabdulla and Hillen (2024) and Al-Tuwairqi et al. (2020). Our model was designed to capture the complex dynamics between cancer cells, oncolytic viruses, and the immune system. The goal of this work was to conduct a detailed qualitative and quantitative analysis of the system in order to address the question: why, despite their promise, do oncolytic viruses as a monotherapy rarely achieve complete and lasting regression of established tumors? We started our work by performing a stability analysis of the space-independent model. This allowed us to identify distinct parameter regimes defined by *θ* (the effective viral production rate) and *η* (the immune stimulation rate by uninfected cancer cells), thus determining the combinations of parameter values under which virotherapy yields the best outcome. We also see in Figure 2 that for large values of *η* the immune system will remove the virus, and the system becomes a pure cancer-immune interaction. For large values of *θ* we observe the opposite, where the immune system is silenced, and we obtain a immune-free dynamics. The combination of oncolytic virus and immune response is most efficient for intermediate values of *η* and *θ*. We continued our parameter analysis with performing a sensitivity analysis of all the parameters of the model. This analysis highlighted the strong influence of the immune-related parameters (*η* and *ζ*) on treatment outcomes. From these findings, we conclude that enhancing the effectiveness of oncolytic virotherapy requires prioritizing strategies that strengthen its dependence on virus-related parameters—particularly the virus’s burst rate *θ*, and the immune stimulation *η* and exhaustion *ξ*.

In the second part of our work, we returned to the reaction–diffusion model to study the spatial dynamics of the system. We identified traveling-wave solutions emerging from the model and, through leading-edge analysis, we found a series of stacked waves that invade at different speeds. Here we like to transform the wave speeds back into the original, dimensional variables. For the cancer invasion front (45), the virus wave (52), and the immune invasion (54), we find

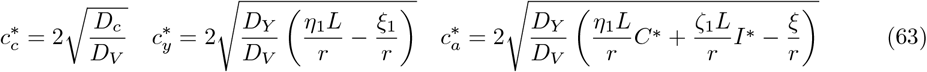

We obtain the speed values in dimensional coordinates in Table 5.

**Table 5:** Waves propagation speeds in dimensional coordinates.

| Case | Wave | Theoretical |
| --- | --- | --- |
| Slow immune | cancer invasion | $c_c = 0.027$ mm/day |
| | virus invasion into the cancer | $c_v = 0.080$ mm/day |
| | immune invasion into cancer | $c_y = 0.028$ mm/day |
| | immune invasion into virus | $c_a = 0.026$ mm/day |
| Fast immune | cancer invasion | $c_c = 0.027$ mm/day |
| | virus invasion into the cancer | $c_v = 0.080$ mm/day |
| | immune invasion into cancer | $c_y = 0.028$ mm/day |
| | immune invasion into virus | $c_a = 0.091$ mm/day |

There is no doubt that many more combinations of wave speeds are possible. We provide explicit formulas for some of the wave speeds in (45), (52), and (54), which directly depend on the model parameters. Once these parameters are known, estimates of these invasion speeds can be made.

Stacked waves suggest that timing matters. If the virus wave outruns the immune wave too much, immune clearance may come too late, risking viral exhaustion or tumor regrowth. Conversely, if the immune wave is too fast, it may wipe out the virus prematurely, which is a common problem in clinical trials of oncolytic virotherapy. Understanding this mathematically helps in treatment scheduling: e.g., spacing doses so that we have the waves overlap more optimally. This is achieved by a more optimized schedule of the virotherapy. While no clinical paper explicitly says “stacked waves,” the phenomenon of multi-phase tumor response (rapid shrinkage followed by slower stabilization or regrowth) matches this idea. In the future, stacked-wave analysis might guide design of combination therapies, optimizing virus dosing schedules that balance the speeds of viral spread and manipulating immune response for maximum tumor eradication.

The stacked waves found here are also mathematically interesting. Similar to the Fisher-KPP equation, travelling waves can be related to heteroclinic orbits in phase space (Vries et al., 2006). A travelling wave analysis of our model (43) would lead to an 8 dimensional phase space. Numerically, we find heteroclinic connections from *E*_1_ to *E*_0_, from *E*_2_ to *E*_1_, from *E*_3_ to *E*_1_, and from *E*_3_ to *E*_2_. Essentially a homoclinic tree as sketched in Figure 10. To prove the existence of such a connected tree of heteroclinic orbits is a formidable challenge, which we cannot solve here.

**Figure 10:**
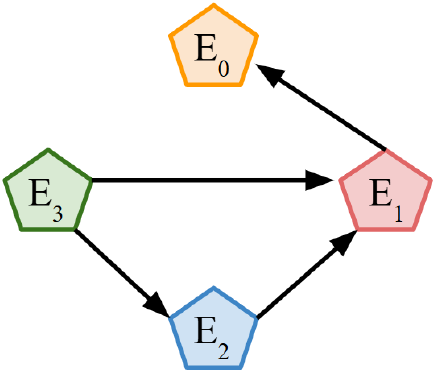
Schematic of the heteroclinic connections between equilibria that comprise the stacked wave. Each of these connections represents an invasion front that might travel at different speeds.

## Acknowledgments

We are particularly grateful to discussions with Morgan van Walsum during her NSERC-USRA 2024 scholarship at the University of Alberta. We also thank the Math Bio Journal Club for helpful comments. NM acknowledges the funding support from the University of Alberta. TH is supported through a Discovery Grant of the Natural Science and Engineering Research Council of Canada (NSERC), RGPIN-2023-04269.

## Data Availability

No data were generated or analyzed during the course of this study, so data sharing is not applicable.

